# Fast calcium-dependent reorientation of motile cilia basal bodies in the simple metazoan, Trichoplax

**DOI:** 10.1101/2025.08.22.671430

**Authors:** Marvin Leria, Moina-mkou Daroueche, Magali Requin, Roman Hill, André Le Bivic, Raphaël Clément, Andrea Pasini

## Abstract

Ciliary-based animal locomotion relies on the spatially coordinated beating of motile cilia to displace the body in liquid environments or across solid surfaces. The orientation of ciliary beating largely depends on the rotational polarity of the basal body, which in most animals is fixed and controlled by cues linked to the main body axes. The small marine animal Trichoplax exploits the beating of motile cilia in its lower epithelium to crawl on substrates. However, Trichoplax lacks defined body axes and exhibits rapid changes in body shape and direction of movement, thus raising the question of what controls the orientation and reorientation of ciliary beating. We show here that the basal bodies of the cilia in the lower epithelium of Trichoplax are oriented along the direction of the animal movements, and that they change their orientation throughout the lower epithelium in a few seconds’ time when the animal modifies its shape or changes direction following external mechanical stimuli. We also show that Ca^2+^ is required for fast basal body reorientation. Such rapid ciliary basal body reorientation has never been observed in metazoans before. Thus, our results identify a previously undescribed mechanism underlying directional motility in a metazoan and shed light on its subcellular determinants, bridging the scale between intra-cellular ciliary machinery and animal movement.

## INTRODUCTION

Most animals rely on locomotion in order to escape threats, migrate to new environments and look for food or partners. Ciliary locomotion, dependent on the directionally coordinated, periodic beating of cilia that generate forces leading to the displacement of the body, is present in a wide range of metazoan phyla, and is likely to be among the most ancient locomotion modes (Marinkovic et al., 2019). The orientation of ciliary beating is determined by interacting physical (e.g. hydrodynamic flow), biochemical (Ca^2+^ and K+ concentration, dopamine and serotonin neurotransmitters, Planar Cell Polarity signals) and structural factors (Mitchell et al., 2007; Guirao et al., 2010; Spassky and Meunier, 2017; Boutin and Kodjabachian, 2019, Wan and Poon, 2023). Among the latter, the rotational polarity of the basal body (BB), the complex multiprotein structure that anchors each motile cilium within the cytoplasm of ciliated cells, plays a particularly important role (Wallingford, 2010, Mirzadeh et al., 2010). The BB generally consists of a cylindrical, centriole-derived structure whose axis extends the ciliary axoneme, and of several asymmetrically attached protruding appendages, among which the most prominent are the basal foot and the striated ciliary rootlet (Mahen, 2021). Basal foot and ciliary rootlet are attached to the centriole at diametrically opposed positions, and define the axis of ciliary beating (Spassky and Meunier, 2017; Boutin and Kodjabachian, 2019).

Although in most cases motile cilia act as tiny oars, allowing adult animals or their larvae to swim within the water column, a smaller number of animals use ciliary beating to ‘crawl’ or ‘glide’ on solid substrates. Among these, the best characterised are Planarians (Platyhelminthes, Tricladida), flatworms which glide on substrates thanks to the beating of thousands of motile cilia covering their ventral surface (Rompolas et al., 2010). Planarians are Bilaterians, and the active stroke of the beating cilia is always directed posteriorwards, so as to propel the animal body forward. The rotational polarity of the BBs (as defined by the rootlet orientation) is fixed and largely aligned along the antero-posterior (AP) body axis, with only minor medio-lateral deviations away from the AP axis around the edges of the body (Vu et al., 2019).

Placozoans are a small phylum of non-bilaterian, millimeter-scale, mostly benthic marine animals with peculiar features that set them apart from all other metazoans. Largely consisting of two layers of monociliated epithelial cells that delimit a closed inner cavity, they lack multicellular organs, muscles or a neuron-based nervous system (Leria et al., 2024). The two best-studied Placozoans, *Trichoplax adhaerens* (Schulze, 1883) and the phylogenetically closely related *Trichoplax* spH2 (Voigt et al., 2004), actively crawl on submerged substrates by means of the beating of the thousands of motile cilia present on the lower epithelium (Smith et al., 2015; Bull et al., 2021a; Bull et al., 2021b). Despite the lack of a nervous system, *T. adhaerens* and *Trichoplax* spH2 move in response to a variety of attractive and repulsive cues (Mayorova et a., 2018; Senatore et al., 2017; Smith et al., 2015; Smith et al., 2019; Ueda et al., 1999; Zhong et al., 2023) but, unlike Planarians, do not show an apparent AP axis and can crawl in any direction. They also display high morphological plasticity, being able to transition from an approximately round, pancake-like shape to branched or filamentous morphologies in a few minutes (Prakash et al., 2021). Neurotransmitters and neuropeptides were shown to induce specific behavioral responses accompanied by body shape changes, locomotion patterns and beating direction of cilia (Senatore et al., 2017; Varoqueaux et al., 2018; Romanova et al., 2020; Nikitin et al., 2023; Jin et al., 2024). It has been recently shown that the direction of ciliary beating can change rapidly upon reorientation of animal crawling (Smith et al., 2015; Smith et al., 2019; Romanova et al., 2020; Bull et al., 2021b) but how this correlates with the orientation of the ciliary polarity machinery is still not known.

Here, using cell-resolved imaging of BBs at the scale of entire animals, we report that the BBs in the lower epithelium of *Trichoplax* spH2 are largely aligned with the global direction of movement of the animal. We also show that the animals change direction in response to mechanical stimuli, and that the BBs reorient accordingly on the timescale of seconds. Besides, we find that Ca^2+^ is necessary for this reorientation to occur. Similar Ca^2+^-dependent fast BB reorientation events have been described in unicellular green algae (McFadden et al., 1987), but never before reported in animals. By bridging the scale from individual cell-level ciliary machinery to whole body-level animal movement, our work sheds light on the cellular determinants of the environment-dependent locomotory behavior in a non-bilaterian animal lacking muscles or a neuronal network-based nervous system.

## RESULTS

### Motility of control Trichoplax

As described for *T. adhaerens*, the lower epithelium of *Trichoplax* spH2 (hereafter Trichoplax for simplicity) (Figure 1A) is covered by thousands of motile cilia (Figure 1B), whose beating propels the animal crawling over the substrate (Movie 1). Trichoplax transferred from algae-coated glass Petri dishes to a clean coverslip quickly adhere to the substrate and perform random movements to explore their environment (Figure 1C; Movie 2). In these conditions, animals typically move over a distance of one body size every minute, with occasional stops (Figure 1D). Given the absence of AP polarity and the morphological plasticity of Trichoplax, these movements can be accompanied by various degrees of body-shape changes (Figure 1E; Movie 3).

**Figure 1.**
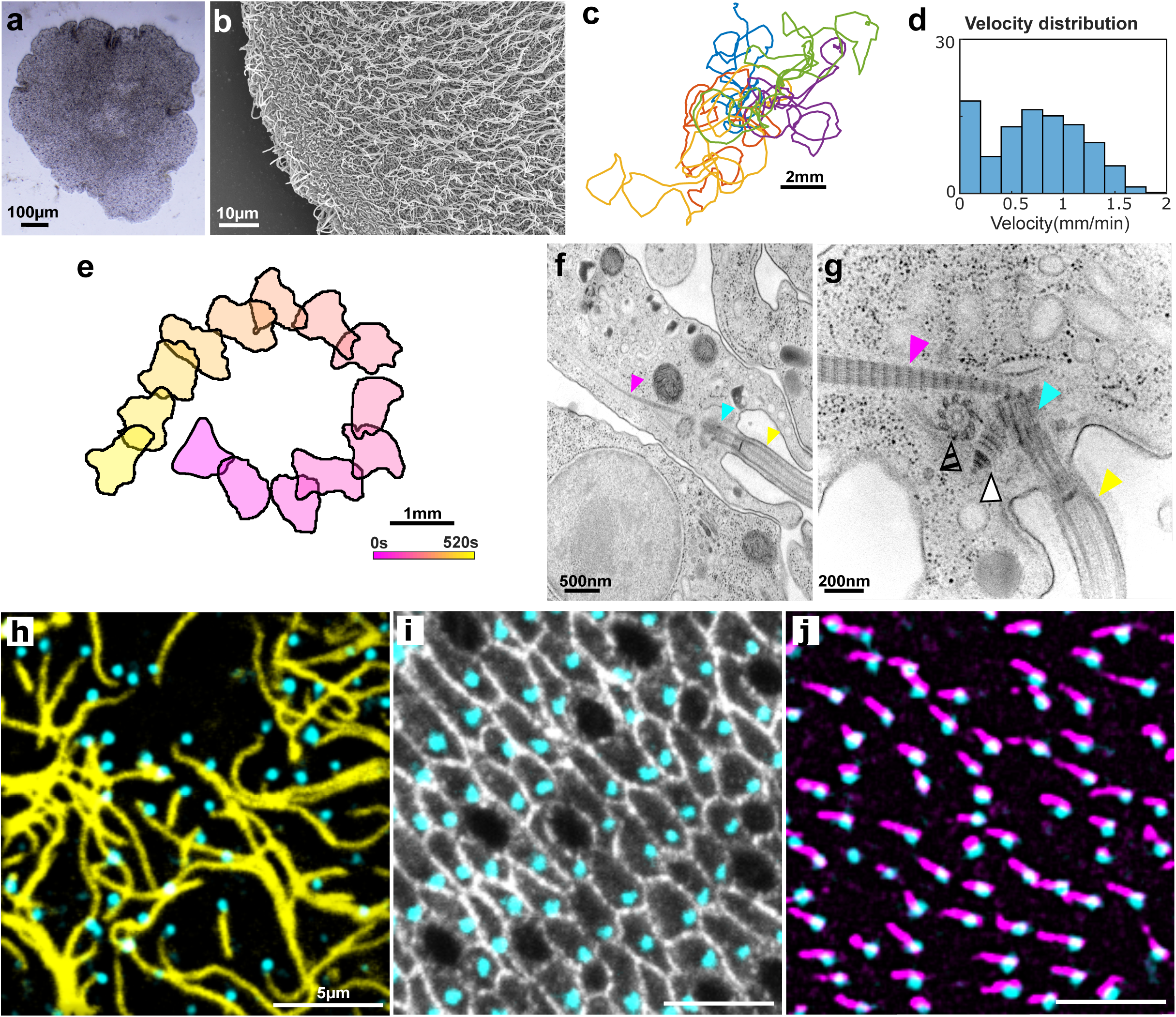
**A:** Brightfield image of a live Trichoplax. **B:** Scanning Electron Microscope image of the lower epithelium of Trichoplax, covered with motile cilia. **C:** Tracks showing the movements of five Trichoplax. **D:** Velocity distribution of moving Trichoplax: each column represents the probability that the animals tracked in C perform a movement at a given speed. **E:** Body outlines of an individual moving Trichoplax (corresponding to the orange track in D) drawn at 40s intervals over a total time of 520s. Body shape changes occur concomitantly with the animal movement. **F:** Transmission Electron Microscopy (TEM) image of a ciliated lower epithelial cell, showing the cilium (yellow arrowhead), the modified basal body centriole (light blue arrowhead) and the ciliary rootlet (magenta arrowhead). **G:** Higher-magnification TEM image of a different ciliated lower epithelial cell, to show the relationships among the different ciliary complex components. Colour codes as in Figure 1F, plus white arrowhead pointing at the basal foot and hatched arrowhead pointing at the sister centriole. **H:** En face confocal microscopy image of the lower epithelium, showing that BB centrioles (light blue, anti-centrin Ab) are each located at the basis of a motile cilium (yellow, anti-Acetylated Tubulin Ab). **I:** En face view of the lower epithelium, showing the apical outlines of lower epithelial cells (white, anti-MAGUK Ab) and the BB centrioles (light blue, anti-centrin Ab). The black cells not showing a centrin signal are the Lipophile cells (Smith et al., 2014). **J:** A view of the lower epithelium showing the BB centrioles (light blue, anti-centrin AB) and the corresponding rootlet (magenta, anti-rootletin Ab).

### Organisation of lower epithelium ciliary basal bodies in Trichoplax

As a first step to explore the correlation between rotational polarity of the ciliary BBs and animal movements, we exploited Transmission Electron Microscopy (TEM) and ImmunoFluorescence (IF) to characterise the lower epithelium BB structures. TEM showed that the ciliary BBs in the lower epithelium are each associated with a striated rootlet extending deep within the cytoplasm (Figure 1F, G). IF with antibodies directed against Acetylated Tubulin (AcTub) to label the ciliary axoneme and against centrin, a major component of the ciliary BB, revealed the presence of a centrin-positive BB at the base of each AcTub-positive cilium on the lower epithelium (Figure 1H). IF with antibodies against centrin and against the Membrane-Associated GUanilate Kinase (MAGUK) proteins to label the cell contours showed that each cell in the lower epithelium, with the exception of the round-looking Lipophile Cells, carries a centrin-positive basal body (Figure 1I). IF with antibodies raised against Trichoplax rootletin and against centrin revealed rootletin-positive elongated structures associated with each BB and compatible with ciliary rootlets (Figure 1J). Altogether, these observations show that the overall structure of the Trichoplax motile cilia BB does not substantially differ from that reported for other Metazoans and that the rootlet can be used as a basal body polarity marker.

### Ciliary basal body orientation is aligned with the direction of movement of freely-moving Trichoplax

To assess the relation between ciliary BB orientation and the direction of movement, individual Trichoplax crawling on glass coverslips were filmed for 20-30 seconds, rapidly fixed, then processed for IF with antibodies against rootletin and centrin and imaged by confocal microscopy (Figure 2A-D). To obtain a high-resolution view of the entire lower epithelium of each individual animal, images taken at single cell-resolution were stitched together (Figure 2A). Using automated segmentation of the centrin and rootletin signals, we defined for each ciliary BB a polarity vector reporting the orientation of the ciliary rootlet and pointing towards the centrin signal (Figure 2E-G). Since Trichoplax lower epithelia typically consists of tens of thousands of cells, polarity vectors were binned into 10µm x 10µm square subregions, each containing approximately 20-25 ciliated cells, to allow better visualization. Thus for each analysed individual (n=5) we obtained a color-coded map of BB orientation across the entire lower epithelium, as well as a polar histogram of BB orientation (Figure 2H). This approach revealed that the average orientation of BBs is aligned with the direction of animal movement immediately before fixation (Figure 2I-L, Movie 4 and Supplementary Figure S1). Remarkably, in individuals that combine movements of their center of mass with significant body shape changes, we observed large-scale gradients of BB orientation, and thus a higher spread of the polarity distribution that fits with the observed body shape changes (Figure 2M-P and Movie 5). Together, these results show that the mean orientation of the BBs points towards the global direction of animal movement, and suggest that gradients of BB orientation, and thus of ciliary beating, can cause in-plane tissue strains eventually leading to animal-scale shape changes.

**Figure 2.**
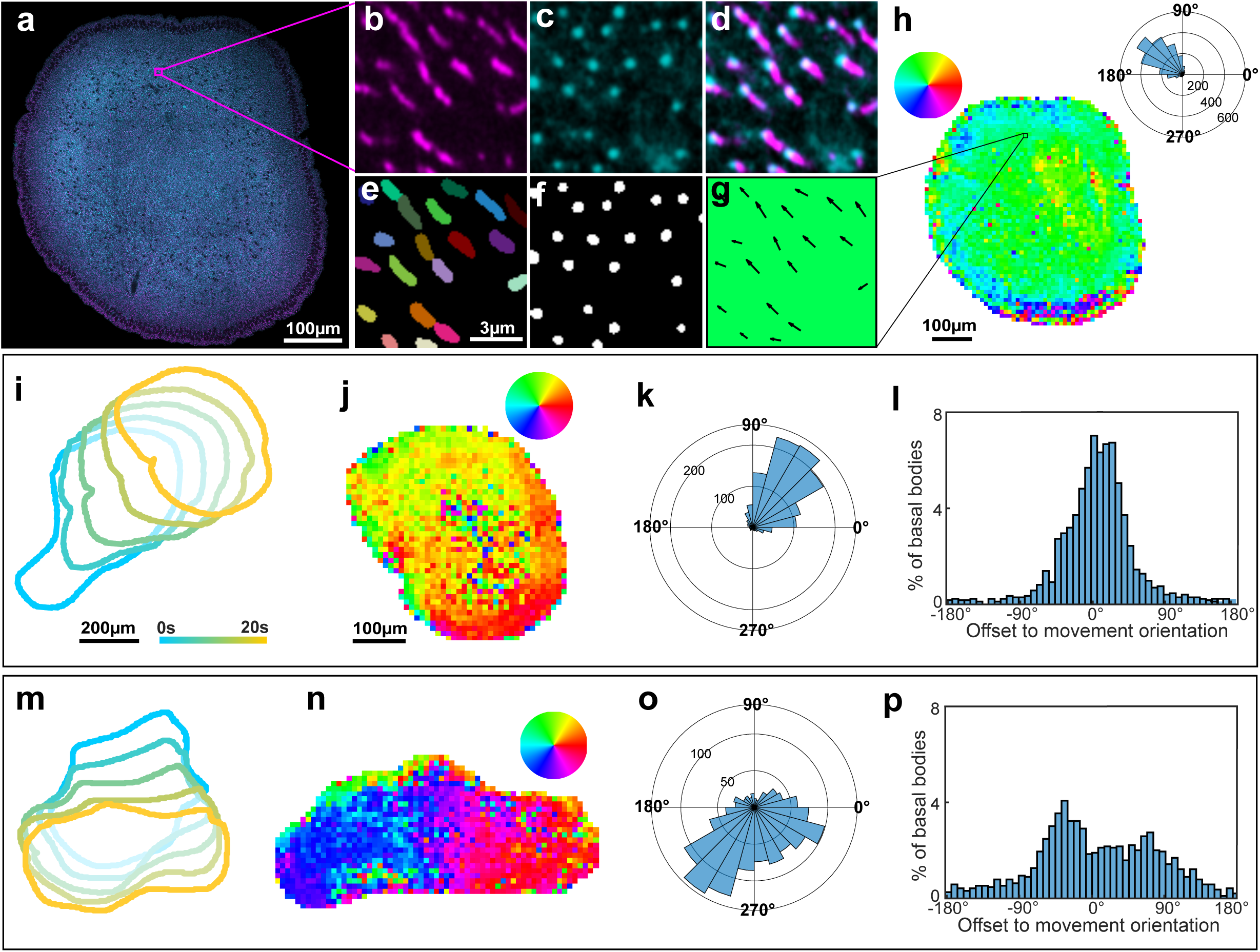
**A:** En face view of the whole lower epithelium of a Trichoplax stained with anti-centrin (light blue) and anti-rootletin (magenta) Abs. Image reconstructed by stitching together nine high resolution confocal images. **B:** A close-up of the boxed area in A, showing the rootletin signal. **C:** A close-up of the same boxed area, showing the centrin signal. **D:** Merge of the centrin and rootletin signals. **E:** Segmentation of the rootletin signal shown in B. **F:** Segmentation of the centrin signal shown in C. **G:** Rotational polarity vectors computed from the signals showed in E and F. **H:** Colour-coded map of BB polarity orientation and polarity histogram for the animal shown in A. The hsv wheel determines the colour code attributed to each trigonometric orientation. **I:** Body outlines of a moving Trichoplax, drawn at 5 second intervals to show the animal displacement. The colour code corresponds to the different times at which the outlines were drawn. **J:** Colour-coded map of BB polarity for the animal in I. **K:** BB polarity histogram for the same animal, showing a global alignment with the direction of movement. **L:** Graph showing the offset of BB orientation respective to the direction of movement of the centroid of the different outlines in I. **M:** As in I, but for a Trichoplax in which the body displacement is combined with significant body shape changes. **N:** Colour-coded map of BB polarity for the animal in M, showing a steep gradient in the polarity orientation. **O:** Basal BB histogram for the same animal, showing a higher spread of the polarity distribution. **P:** Graph showing the wide offset of BB orientation respective to the direction of movement of the centroid of the different outlines in M.

### Fast reorientation of animal movements and basal body polarity following mechanical stimuli

Chemo-and thermotactic cues are known to affect the directionality of Trichoplax movements (Senatore et al., 2017; Smith et al., 2015; Smith et al., 2019; Romanova et al., 2020; Varoqueaux et al., 2018; Zhong et al., 2023) but the role of mechanical stimulation has never been thoroughly explored. We thus set out to perform mechanical stimulation on Trichoplax with a two-fold aim: to assess whether this can modify the animal movements, and to determine its possible impact on the orientation of the lower epithelium BBs. Two types of stimuli were applied: poking and bisection. In poking experiments, the leading edge of freely-moving individuals (n=10) was gently touched with a round-tip microscalpel. This resulted in the animals reorienting their movement in a direction on average opposite to that of the stimulus after a few seconds (Figure 3A,B and Movie6). This reorientation occurred a few seconds after the stimulus and persisted for at least 30 seconds, showing that the animals were not simply pushed away by the touch, but actively crawled away from it. To determine whether this active movement reorientation was correlated with a repolarisation of the BBs, animals (n=3) were fixed 30 seconds after the stimulation, stained with antibodies against centrin and rootletin and analysed as described in the previous paragraph. Interestingly, this revealed that 30 seconds after applying the mechanical stimulus, the majority of BBs were oriented in the new direction of movement (Figure 3C, D and Supplementary Figure S2 A and B).

**Figure 3.**
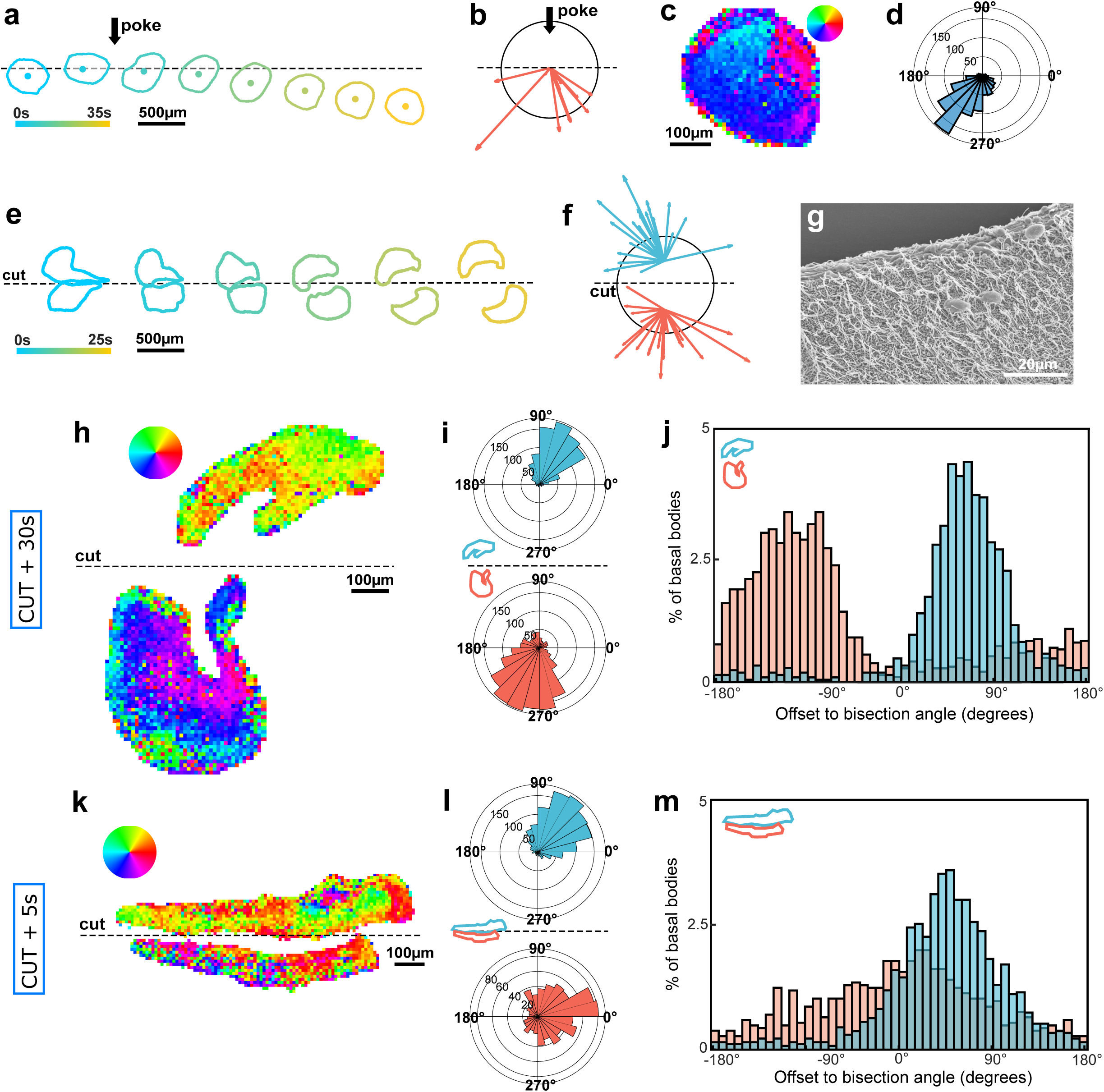
**A:** Body outlines of a moving Trichoplax, before and after poking (black arrow), drawn at 5 second intervals. The colour code corresponds to the different times at which the outlines were drawn and the centroid of each outline is indicated. **B:** Centroid displacement for 10 independent poked Trichoplax. The black circle represents an approximation of the individual body outlines and the coloured arrows show the individual centroid displacements. Black arrow: point and direction of poking. **C:** Colour-coded map of BB polarity for the animal in A. **D:** BB polarity histogram for the same animal, showing a global alignment with the direction of centroid displacement after poking. **E:** Body outlines of a manually sectioned Trichoplax. Colour code as in A. **F:** Graph showing the centroid displacement for the two halves of 24 independent bisected Trichoplax. The black circle represents an approximation of the individual body outlines before cutting, the stippled line shows the cutting axis and the coloured arrows the individual centroid displacements. **G:** Scanning Electron Microscope image of the lower epithelium of a bisected Trichoplax showing the motile cilia oriented away from the cutting site. **H:** Colour-coded map of BB polarity for a bisected animal fixed 30 seconds after cutting, showing opposite polarity orientations in the two animal halves. **I:** Basal BB histogram for the two halves of the animal in H. **J:** Offset of BB orientation for the two animal halves in H, respective to the cut. **K:** Colour-coded map of BB polarity for a bisected animal fixed 5 seconds after cutting, showing incomplete reorientation in the two animal halves. **L:** BB histogram for the two halves of the animal in K. **M:** Offset of BB orientation for the two animal halves in K, respective to the cut.

In bisection experiments, Trichoplax (n=24) were manually bisected with a microscalpel. This systematically resulted in the two halves of the animal crawling away from each other, approximately perpendicular to the bisection axis (Figure 3E, F and Movie 7). Analysis of the BB orientation (n=3) revealed that 30 seconds after bisection, the BBs in each of the two animal halves had already reoriented away from the cutting plane and pointed towards the new directions of movement (Figure 3H-J, Movie 8 and Supplementary Figure S2 C and D). Interestingly, in animals immediately (3-5 seconds) fixed after bisection (n=2), the BBs in the two separate halves were found to be still largely oriented along the direction of movement followed by the intact animal, suggesting a minimum time required to perform the coordinated reorientation (Figure 3K-M, Movie 8 and Supplementary Figure S2 E).

Together, these results show that Trichoplax can change its crawling direction in response to mechanical signals, and that this reorientation correlates with a repolarisation of the motile cilia BBs which takes place across the entire lower epithelium on the timescale of seconds.

### Effect of Ca2+ depletion on Trichoplax movement and basal bodies orientation

Our data revealed a phenomenon of BB reorientation taking place across the entire lower epithelium in response to local mechanical stimuli, suggesting the long-range propagation of a reorientation signal starting from the stimulation point. Given the rapidity of reorientation we reasoned that Planar Cell Polarity signalling is unlikely to play a role and instead set out to address the possible role of Ca^2+^. Indeed, propagating Ca^2+^ waves play a role in several instances of coordinated multicellular responses, and across evolution Ca^2+^ signaling controls a number of cellular responses to external cues (Clapham, 2007), one remarkably interesting example being the fast reorientation of ciliary BBs in unicellular green algae (Mcfadden et al., 1987).

Trichoplax were incubated for 1hr in 50mM EGTA, a known chelator of Ca^2+^ ions. Tracking showed that under these conditions, animals (n=6) are viable and motile, although their velocity and the speed of ciliary beating were significantly decreased compared to controls (Figure 4A, B, Movies 9 and 10). IF with antibodies against centrin and rootletin followed by analysis of the lower epithelium ciliary BB orientation (n=3) revealed an overall alignment with the direction of animal movement (Figure 4C-E and Movie 11). In contrast, following bisection of EGTA-treated Trichoplax (n=22), the two animal halves did not significantly move away from each other (Figure 4F-H and Movie 12) and the lower epithelium BBs failed to reorient away from the bisection plane as observed in cut, untreated controls. Instead, they either showed the same orientation in the two animal halves or random orientation (n=3) (Figure 4I, J).

**Figure 4.**
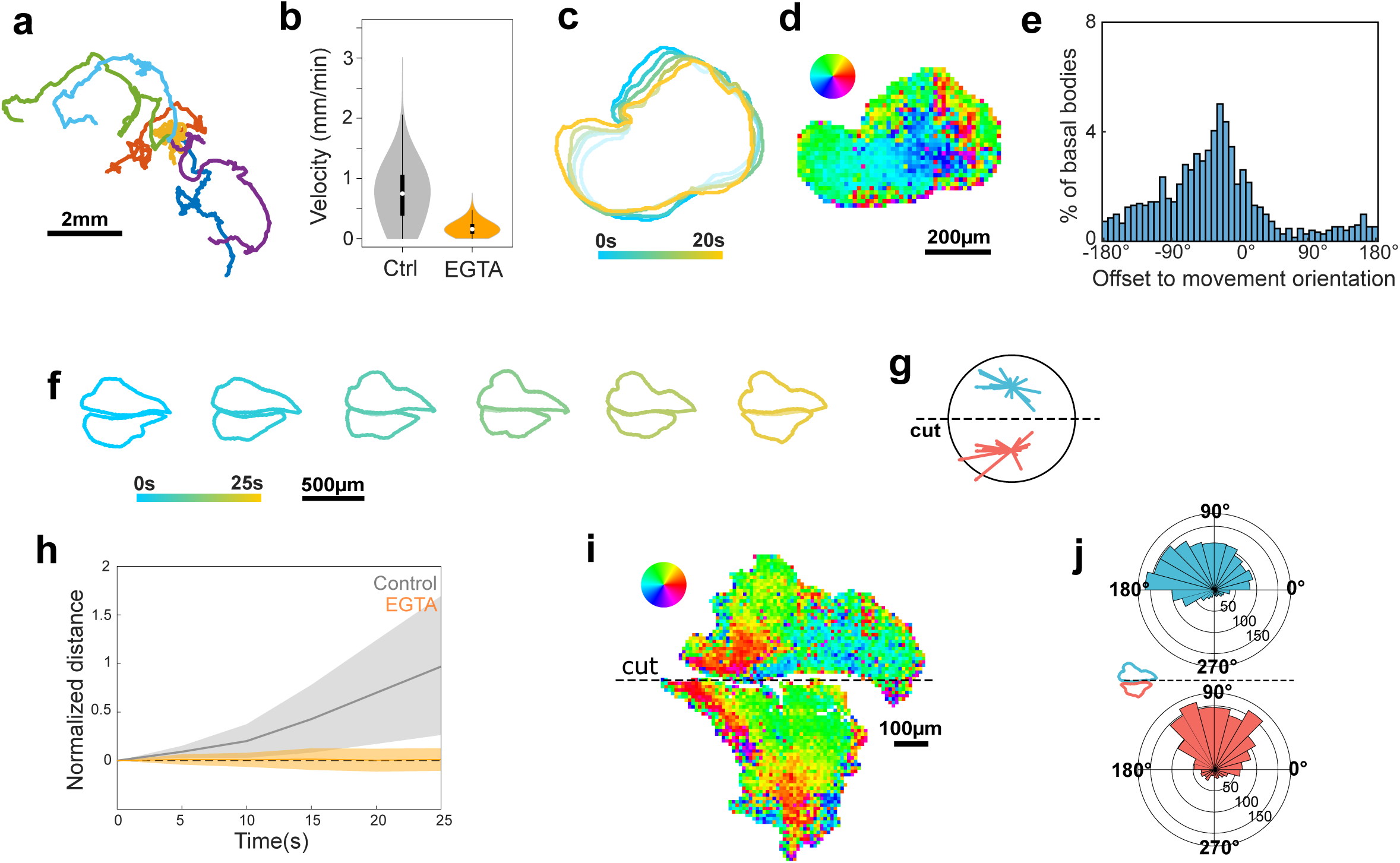
**A:** Tracks showing the movements of six Trichoplax incubated over 1hr in 50mM EGTA. **B:** Speed distribution of moving Trichoplax in 50mM EGTA. **C:** Body outlines of a moving Trichoplax in 50mM EGTA, drawn at 5 second intervals. **D, E:** Colour-coded map of BB polarity and offset of BB orientation for the animal in C. **F:** Body outlines of a sectioned EGTA-treated Trichoplax. Colour code as in C. **G:** Centroid displacement for the two halves of 22 bisected EGTA-treated Trichoplax. **H:** Graph showing the mutual distancing of the halves of bisected control (black) and EGTA-treated (orange) animals. The grey and light orange areas represent the standard errors for the two conditions. **I, J:** BB polarity map and BB histogram for a bisected EGTA-treated animal fixed 30 seconds after cutting, showing lack of reorientation in the two animal halves.

Thus, our results show that the environmental depletion of free Ca^2+^ ions perturbs the reorientation of BBs following Trichoplax bisection, suggesting that Ca^2+^ is required for this phenomenon.

### Pharmacological inhibition of Voltage-Gated Calcium Channels affects Trichoplax movement and basal body reorientation

To further characterize the role of Ca^2+^ signalling in BB reorientation, we treated Trichoplax with Verapamil, a pharmacological inhibitor of Voltage-Gated Calcium Channels (VGCCs) (Zhao et al., 2019). VGCCs, which allow Ca^2+^ influx into the cells following membrane depolarization, are crucial actors in triggering fast cellular responses to external stimuli and have been identified in Trichoplax (Senatore et al., 2016). Tracking of animals (n=5) treated for 1hr with 75µM Verapamil revealed a slowing down of their overall animal velocity compared to untreated controls (Figure 5A, B; Movie 13). However, the lower epithelium BBs were globally oriented in agreement with the direction of animal movement (n=1) (Figure 5C-E and Movie 14). On the other hand, and similar to what observed following EGTA treatment, in Verapamil-treated Trichoplax, bisection (n=20) did not lead to a significant distancing of the two animal halves (Figure F, G and Movie 15) and BBs failed to reorient away from the cut, showing instead persistent or random orientations (n=2) (Figure 5H-J). Our data thus uncover a role for VGCC-mediated influx of Ca^2^ in BB reorientation after bisection.

**Figure 5.**
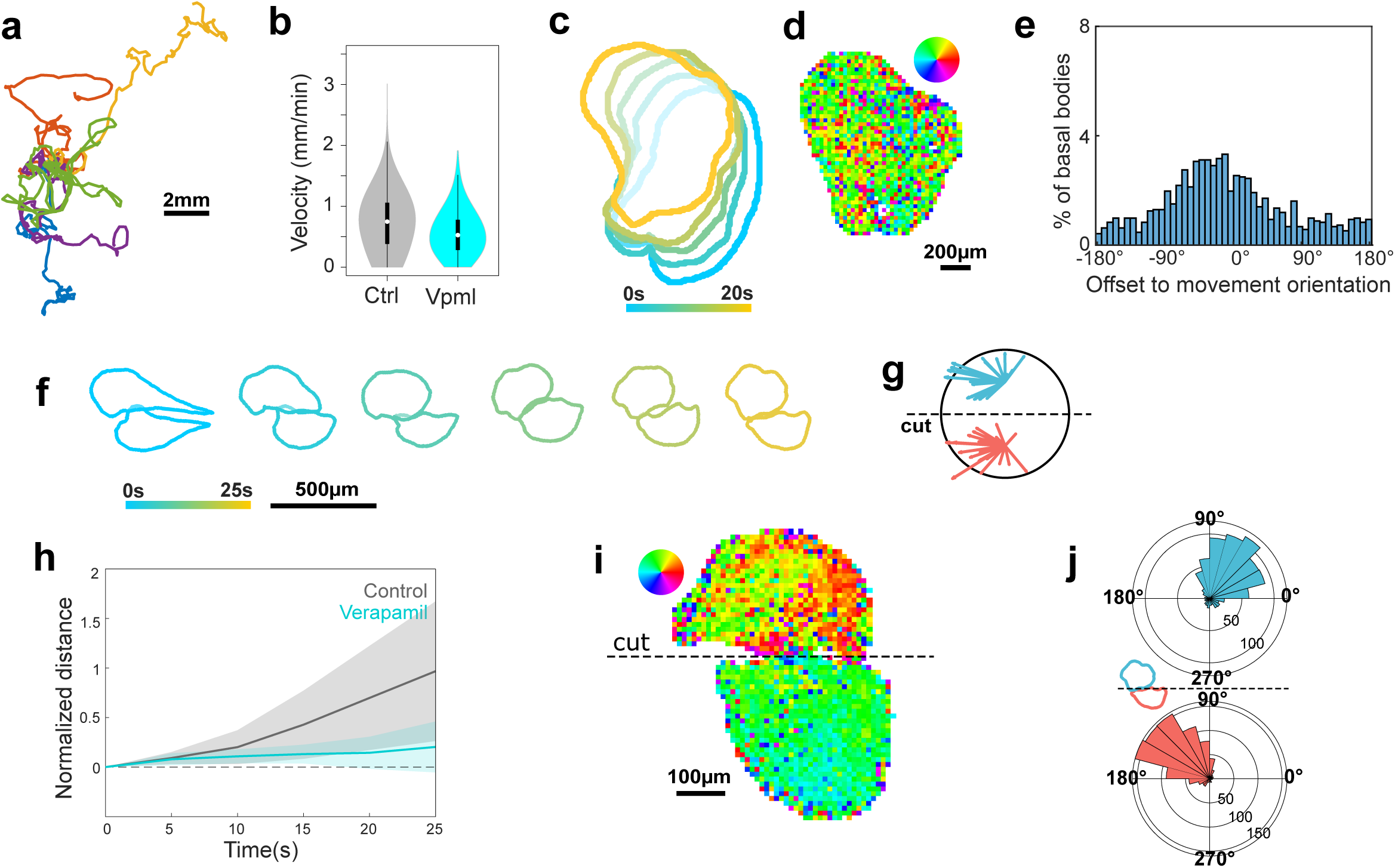
**A:** Tracks showing the movements of five Trichoplax incubated over 1hr in 75µM Verapamil. **B:** Speed distribution of moving Trichoplax in 75µM Verapamil. **C:** Body outlines of a moving Trichoplax in 75µM Verapamil, drawn at 5 second intervals. **D, E:** Colour-coded map of BB polarity and offset of BB orientation for the animal in C. **F:** Body outlines of a sectioned Verapamil-treated Trichoplax. Colour code as in C. **G:** Centroid displacement for the two halves of 20 bisected Verapamil-treated Trichoplax. **H:** Graph showing the mutual distancing of the halves of bisected control (black) and Verapamil-treated (blue) animals. Light grey and light blue areas: standard errors for the two conditions. **I, J:** BB polarity map and BB histogram for a bisected Verapamil-treated animal fixed 30 seconds after cutting, showing lack of reorientation in the two animal halves.

## DISCUSSION

Locomotion confers animals the ability to change their immediate environment, typically to find food or escape threats. Studying the cellular basis of locomotion in early diverging phyla is key to understanding the emergence of the locomotory behaviour in metazoans and its evolutionary diversification.

Ciliary locomotion is an evolutionarily ancient mode of locomotion, exploited by many uni-and multicellular organisms across several phyla (Marinkovic et al., 2019) and relying on the spatially-coordinated beating of motile cilia to achieve motion within liquids (ciliary swimming) or over solid surfaces (ciliary gliding). The beating orientation of motile cilia depends on both extrinsic (e.g., environmental fluid flows) and intrinsic (e.g., cell structural or biochemical factors) cues (Mitchell et al., 2007; Guirao et al., 2010; Spassky and Meunier, 2017; Boutin and Kodjabachian, 2019). The rotational orientation of the BB, the multiprotein structure which anchors the motile cilia within the cytoplasm, plays a major role in defining the orientation of ciliary beating (Wallingford, 2010). In most metazoans, the BB orientation is set up during embryonic development, is aligned along the main body axes and is largely maintained throughout life (the most notable exceptions being metamorphosis and some regenerative processes). The Planar Cell Polarity (PCP) pathway as well as the actin and tubulin cytoskeletal components and molecules that link them to the BB have been variously involved in setting up rotational polarity in several species.

Here, we examine the curious case of Trichoplax, an anatomically simple, morphologically plastic non-bilaterian marine benthic animal, which lacks major body axes and moves by ciliary walking while constantly changing body shape. Recent work has shown that the cilia on the lower epithelium of Trichoplax can coordinately change their beating direction in a few seconds, thus allowing the animal to modify its direction of locomotion (Smith et al., 2015; Smith et al., 2019; Romanova et al., 2020; Bull et al., 2021b).

The molecular and structural underpinnings of ciliary beating reorientation in Trichoplax have never been addressed. Our results show that the predominant orientation of the BBs of the lower epithelium motile cilia corresponds to the overall direction of animal locomotion. Interestingly, in individuals simultaneously undergoing locomotion and body shape changes, we observed BB orientation gradients, which are likely to generate mismatches in crawling direction. The resulting in-plane strains can lead to whole-body shape changes, but are also likely to underlie more elaborate movements, such as fold flattening (Brannon and Prakash, 2024) or spontaneous epithelial fractures (Prakash et al, 2021). How such gradients appear is unclear, but they might be the outcome of biochemical signals and/or stem from a trade-off between body size and spatial coordination (Davidescu et al., 2023). These observations suggest a phenomenon of ciliary BB planar reorientation underpinning the fast and coordinated changes in the beating direction of the lower epithelium cilia that allow Trichoplax to perform intricate movements and shape changes on the timescale of seconds.

Accordingly, we found that mechanical stimuli, such as large wounds or more localised ‘poking’, systematically result in a fast (second-timescale) reorientation of BBs across the lower epithelium and in the reorientation of animal movement. A very tempting hypothesis, to be further investigated, is that such a fast reorientation mechanism in response to mechanical stimuli might represent a successful strategy to sense obstacles and/or escape noxious mechanical stimuli that emerged and was selected in Placozoans, which lack both muscle and a nervous system based on interconnected neurons.

To our knowledge, such rapid BB reorientation has never been described before in metazoans, and could be exclusive to the morphologically plastic Placozoans. Indeed, the few reported cases of BB planar reorientation in animals occur either in the context of embryonic development, as in Xenopus MultiCiliated Cells (MCCs) antero-posterior polarisation (Boutin and Kodjabachian, 2019), sea urchin and jellyfish monociliated cells (Mizuno et al., 2017; Momose et al., 2012) or during regenerative processes, as described for the repolarisation of MCCs in regenerating planarians (Vu et al., 2019). In both cases, reorientation takes hours to days to complete. In contrast, a similarly rapid BB reorientation has been observed (although in a largely different geometric configuration) in the green alga *Spermatozopis similis*, in which it depends on Ca^2+^ influx into the cell (McFadden et al., 1987), and has been hypothesized for sea urchin larvae (Wada et al., 1997).

Exploring the possible role of Ca^2+^, we found that environmental Ca^2+^ and VGCC-mediated cellular Ca^2+^ influx cells are required for fast BB reorientation following mechanical stimuli in Trichoplax. Ca^2+^, alone or in combination with monoamine neurotransmitters such as dopamine, serotonin or epinephrine, has already been shown to directly affect the ciliary beating frequency, direction and coordination in a variety of bilaterian and non-bilaterian adult marine animals or in their larvae, including *Trichoplax adhaerens* (Tamm and Tamm, 1981; Nakamura and Tamm, 1985; Moss and Tamm, 1987; Wada et al., 1997; Tamm and Terasaki, 1994; Wong et al., 2022; Jin et al., 2024). However, in almost all the reported cases, the observed changes in beating orientation consist in a simple 180° reversal, which leads to the organisms swimming backwards, and are not associated with BB rotational reorientation. A likely scenario to explain our results is that Ca^2+^ influx directly affects the rotation of BBs by acting on structural proteins and that this entails a rotation of the entire ciliary complex. Several possible Ca^2+^-dependent structural determinants of BB rotation can be envisaged and their actual role in this process will have to be further determined. Actin and tubulin cytoskeletal components, known to be important in determining and stabilising the position of BBs in mono-and multiciliated cells (Werner et al., 2011; Antoniades et al., 2014; Donati et al., 2021), might also play a role in the fast reorientation phenomenon described here. Indeed, an involvement of the actomyosin cytoskeleton has been recently postulated in similarly fast cellular events occurring in Trichoplax (Armon et al., 2018). The BB-associated protein centrin can undergo conformational changes upon Ca^2+^ binding, and centrin or centrin-like proteins have been involved in extremely fast, Ca^2+^-induced structural changes in several unicellular organisms (McFadden et al., 1987; Upadhyaya et al., 2008; Lannan et al., 2024; Qin et al., 2024). Finally, Ca^2+^-dependent contractility of ciliary rootlets themselves, resulting in BB displacement has been described in unicellular flagellate algae (Salisbury and Floyd, 1978).

Trichoplax lower ciliated epithelium being composed of tens of thousands of monociliated cells, the coordinated response to a local mechanical stimulation requires the propagation of the BB reorientation signal across the entire epithelium. It is tempting to speculate that Ca^2+^, notably in the form of Ca^2+^ waves, is also involved in such a long-range process. Indeed, tissue-wide Ca^2+^ waves in response to mechanical stimulation are a common feature across evolution (Razzell et al., 2013; Restrepo and Basler, 2016; Hagihara et al., 2022; Hashimura et al., 2022; Colgren and Burkhardt, 2025; Zhou et al., 2025) and their propagation speed matches the timescale of BB reorientation described here (>5 seconds, <30 seconds). Trichoplax lack gap junctions, which constitute a widespread strategy of Ca^2+^ wave intercellular propagation, thus alternative hypothetical mechanisms of propagation will have to be envisaged and explored. These might imply the release of secreted molecules such as NO, monoamine neurotransmitters and soluble neuropeptides, as well as the direct transmission of mechanical or electrical signals, as observed in glass sponges and choanoflagellates (Leys et al., 1999, Colgren and Burkhardt, 2025).

Altogether, this work opens new avenues to understand the cellular and molecular bases of an environment-dependent locomotion behavior in an early-diverging phylum. It also raises a number of questions on the sensing mechanisms exploited by a neuronless animal to probe its environment, to modify accordingly the polarity of its ciliary structures, and to eventually implement various locomotory behaviors. In the absence of a neuron-based nervous system, the variety of behaviors observed in Trichoplax, such as diffusive motion, shape changes, fission, taxis, or fold flattening, all have to stem from cellular feedback rules and propagation mechanisms, making this animal an ideal system to study self-organized, emergent coordinated locomotion.

## MATERIALS AND METHODS

### Animal collection and maintenance

Trichoplax were collected from racks of glass microscope slides that were placed on the bottom of the aquariums of a local pet store in Marseille, France. After three weeks, the slides were recovered, transferred to the laboratory and observed with a binocular dissection microscope. Placozoans were identified, retrieved and transferred by pipetting to large glass Petri dishes containing 100ml filtered artificial sea water (FASW) made by dissolving 33g/L Instant Ocean Sea Salt (#218035, Aquarium Systems, Sarrebourg, France) in milliQ H20, where they were maintained at 23°C under a 12hrs/12hrs dark-light cycle. Animals were fed weekly a mix of *Rhodomonas salina* (gifts from Adriano Senatore, University of Toronto Mississauga, Canada and Harald Gruber-Vodicka, University of Kiel, Germany), *Nannochloropsis oculata* (#T0055, Teramer, Montpellier, France) and *Dunaliella salina* (#T0057, Teramer, Montpellier, France). Water was changed weekly.

For genotyping, 1 and 10 animals were lysed and amplification of mitochondrial *16S rDNA* gene was performed by PCR using the primers and conditions described by Signorovitch et al (Signorovitch et al., 2006). DNA sequences obtained from PCR fragments were BLASTed against the NCBI database resulting in a match with the Trichoplax H2 haplotype.

### Trichoplax movement recording

For recording movements, individual Trichoplax were placed in a drop of FASW on round glass coverslips, left to settle and adhere for a few minutes, then filmed for variable amounts of time under a Leica S4E stereomicroscope equipped with a DinoEye camera or under a Leica S9i stereomicroscope with an integrated camera. Tracking was performed by manually contouring animal outlines every 20 seconds in FIJI.

### Scanning and Transmission Electron Microscopy

For Scanning Electron Microscopy, Trichoplax were placed in a 100ul drop of FASW on glass coverslips, left to adhere, then fixed with Osmium Fixative (NaCl 460mM, Na Cacodylate 100mM, OsO4 78mM, Glutaraldehyde 0,3% in milliQ H2O) for 20min on ice. After rinsing in FASW, samples were postfixed in 0,3% Glutaraldehyde in FASW overnight at 4°C, rinsed again in milliQ H2O, dehydrated by incubation in progressively increasing EtOH concentrations (from 25% v/v in milliQ H2O to absolute EtOH), washed three times with Hexamethyldisilazane, then left to dry overnight. Samples were then sputter-coated with gold using a Edwards S150B Sputter Coater and observed with a FEI Teneo^TM^ VS Scanning Electron Microscope (5,00 kV, spotsize 6-7, dwell time 1-3us, High Vacuum).

For Transmission Electron Microscopy, individual Trichoplax were transferred to 1,4mm diameter, 0,05mm thick, sapphire coverslips (#16706849, Leica Microsystems), left to adhere, submitted to High-Pressure Freezing fixation on a LEICA EMPACT-2 HPF machine, then cryosubstituted with (2% (v/v) OsO4 in Acetone) in a Leica AFS2 Freeze Substitution and Low Temperature Embedding System using the following program:-90 °C for 24 hours;-90°C =>-60 °C over 12 hours;-60 °C for 8 hours;-60°C =>-30 °C over 12 hours;-30 °C for 8 hours; – 30°C =>-5 °C over 1 hour and 15min. At the end of the cryosubstitution process, the samples were washed with Acetone, embedded in 100% Hard Plus Resin-812, cooked at 60°C for 48hrs then cut in 90nm-thick sections with a Leica EM UC7 Ultramicrotome. Sections were contrasted by incubation for 5min in Lead Citrate Trihydrate 3% in milliQ H2O, washed in milliQ H2O and observed on a Tecnai G2 Transmission Electron Microscope running at 200kV.

### Mechanical stimulation and pharmacological treatments

Trichoplax were deposited on round glass coverslips as described above, except that the coverslips were placed within small custom-made holders to facilitate their subsequent handling. For bisection experiments, animals were cut in two halves with a hand-held microscalpel mounted on a pin holder (Phymep, France) then fixed (see below for details) immediately after bisection (within 4s) or after 30s. ‘Poking’ experiments were performed by gently touching the rim of an animal with a hand-held round-tip microscalpel mounted on a pin holder (Phymep, France), followed by fixation 30s after.

For calcium signaling inhibition experiments, EGTA 0.5M pH8 was added to a final concentration of 50mM in seawater for 1hr. To block Voltage-Gated Calcium Channels, Verapamil (REF) was added to a final concentration of 75µM in seawater for 1hr.

### High-speed live imaging of ciliary beating

Individual Trichoplax were deposited on a glass coverslip in a drop of FASW or EGTA-supplemented FASW, left to adhere, covered with a second coverslip using adhesive-tape spacers to prevent squeezing, then observed and filmed in brightfield on a Zeiss Axio M2 microscope, using a Freefly Ember S5K camera (acquisition speed 500 images/s). Movies were treated with Blackmagic’s DaVinci Resolve (https://www.blackmagicdesign.com/products/davinciresolve) to adjust the color temperatures, increase contrast, selecting high-detail frequencies to highlight the cilia and low-detail frequencies to smoothen the contrast of the background epithelial cells.

### Antibodies

The following commercial antibodies were used: mouse monoclonal against centrin (#04-1624, Sigma-Aldrich); mouse monoclonal against acetylated alpha-Tubulin (#T6793, Sigma-Aldrich); mouse monoclonal against pan-MAGUKs (#MABN72, Sigma-Aldrich).

Polyclonal antibodies against Trichoplax rootletin were generated by Eurogentec (Eurogentec SA, Seraing, Belgium) by immunising guinea pigs with the two peptides-CTNYSSPLKSKYGGRT and-CKSPSKVSPRRQTKYK identified in the predicted protein sequence of Trichoplax spH2 rootletin (Uniprot accession number:

A0A369RWE8) (Eitel et al 2018). Antibodies were affinity purified and diluted to a final concentration of 50% glycerol.

### Immunofluorescence staining

For cilia staining, the glass coverslips with adherent Trichoplax were transferred to 3.7% formaldehyde in FASW for 1h at room temperature (RT), then washed 3 times in PBS 1X and permeabilized for 20min in PBS containing 0.1% Triton-X100. Samples were transferred to blocking buffer (goat serum 3%, donkey serum 3%, bovine serum albumin 1% in PBS) for 30min to 1hr then incubated overnight at 4°C with anti-acetylated alpha-tubulin (dilution 1:200) and anti-centrin (dilution 1:1000) antibodies. Samples were washed 3 times in PBS 1X for 10min and were then incubated in blocking buffer with secondary antibody goat anti-mouse IgG2a Alexa Fluor 647 (A-21241, Invitrogen) and goat anti-mouse IgG2b 488 (A-21141, Invitrogen). They were washed 3 times in PBS for 10min and mounted with Prolong Gold antifade reagent with DAPI (P36935, Life Technologies).

For BBs and rootlets staining, we adapted a previously described freeze-substitution protocol (Smith et al., 2014; Smith et al., 2021). Trichoplax on glass coverslips were plunged into cold acetone chilled on dry-ice and stored overnight at-80°C. Samples were then transferred to a mixture of formaldehyde 3.7% in MetOH for 3hrs at-20°C, they were subsequently incubation 5-10 min at-20°C in 2% trichloroacetic acid in MetOH to improve rootletin staining. Samples were rehydrated through downgrading baths of EtOH (from absolute EtOH to 25% EtOH in PBS 1X) every 10min, washed 3 times in PBS 0.1% Triton X-100 for 10min. After blocking in blocking buffer for 1hr, the samples were incubated with a mix of the two anti-Trichoplax rootletin antibodies (each diluted 1:200) and the anti-centrin antibody (1:1000) in blocking buffer overnight at 4°C. They were rinsed 3 times every 10min with PBS 1X and then incubated 3-4h at room temperature or overnight at 4°C in blocking buffer with anti-guinea pig AlexaFluor 488-conjugated antibodies (#706 545 148, JIR) and AlexaFluor 647-conjugated anti-mouse antibodies (#715 605 151, JIR). The same protocol, but replacing acetone with tetrahydrofuran, was used to stain cell outlines with anti-pan MAGUK antibody diluted at 1:500.

### Confocal microscopy

For visualisation of rootletin and centrin staining across the whole lower epithelium, Trichoplax were observed with a Zeiss LSM 880 Airyscan confocal microscope equipped with a 40x water objective using the Tile Scan function with 10% overlap, and with short z-stack, bidirectional line scanning, average 4, laser scanning speed 7, sampling 2140×2140, and simultaneous acquisition of two channels (488 and 633). All other images were acquired with the same confocal microscope or with a Zeiss LSM 780, with objectives 63x oil or 40x water. All images were subsequently analyzed with FIJI.

### Polarity analysis

A dedicated image analysis pipeline was developed to perform BB polarity analysis. For each tiled image, the z-stacks were individually extracted and saved as TIFF files (https://gist.github.com/lacan/16e12482b52f539795e49cb2122060cc), then stitched together using the Grid/Collection Stitching function in FIJI (Preibisch et al., 2009). Maximum intensity projection was performed to optimize signal intensity.

A model was trained in ilastik (Berg et al., 2019) to recognize the centrin signal corresponding to the BB. The centrin signals from all the images were segmented using the ilastik plugin (https://github.com/ilastik/ilastik4ij#ilastik-imagej-modules) in FIJI, the obtained probability map was thresholded and centroid xy-coordinates extracted using the Analyse Particle function. For rootletin signal detection, another model was trained in Cellpose (Stringer et al., 2021) based on the pre-trained CP model. The rootletin signals from all the images were segmented using Cellpose through the FIJI BIOP plugin (https://github.com/BIOP/ijl-utilities-wrappers). ROIs were obtained from the segmentation, fitted to an ellipse and the centroid xy-coordinates, the major axis length and the major axis angle were extracted with the Measure function.

For each BB, the polarity was determined by computing a vector going from the centroid of the rootletin signal to its extremity closest to the centrin signal. The lower epithelial surface of the whole animal was then subdivided into 10µm x 10µm square subregions, each containing around 20-25 ciliated cells and in which the BB polarity vector was averaged, allowing to extract an angle for each square subregion. The polarity orientation of each box was mapped by attributing a colour-code determined by a hsv wheel around the trigonometric circle. All plots were made in MATLAB.

## ACKNOWLEDGEMENTS

This work was supported by doctoral fellowships to M.L from Turing Centre for Living Systems and from the Fondation ARC pour la recherche sur le cancer (grant n°ARCDOC42024010007739). M.R., A.L-B., R.C. and A.P. are CNRS staff members.

This project received funding from the CNRS (‘Diversity of Biological Mechanisms’ 2022 and 2024 to A.P.), from the French Agence Nationale de la Recherche (ANR-21-CE13-0013-01 to A.L-B.) and from France 2030, the French Government program managed by the French National Research Agency (ANR-16-CONV-0001) and from Excellence Initiative of Aix-Marseille University - A*MIDEX. We thank the imaging facility at IBDM, member of the National Infrastructure France-BioImaging (https://ror.org/01y7vt929) supported by the French National Research Agency (ANR-24-INBS-0005 FBI BIOGEN). We thank the owner of the ‘Angel Tropical’ pet shop in Marseilles for allowing us to recover Trichoplax from his aquaria, B. Detailleur for manufacturing racks and holders required for Trichoplax manipulation. We are grateful to R. Kelly and his lab for granting us access to their stereomicroscope S9i, to F. Richard, N. Brouilly and A. Aouane for help with Transmission and Scanning Electron Microscopy, to A. Senatore for providing *R. salina* samples. We thank J. Azimzadeh for her valuable and constructive feedback, B. Charrier and L. Guignard for insights in the project and past and present members of the Le Bivic laboratory for discussions, help and technical support. A.P. especially thanks B. Schierwater and members of his laboratory for introducing him to Trichoplax.

## AUTHOR CONTRIBUTIONS

Conceptualization: M.L, R.C and A.P.; methodology: M.L., M.D., M.R., R.H., R.C. and A.P.; software: M.L. and R.C.; formal analysis: M.L., M.D. and R.C.; resources: A.L-B.; supervision: R.C. and A.P.; project administration: A.L-B. and A.P.; funding acquisition: A.L-B. and A.P.; writing: M.L., R.C. and A.P.

## MOVIES

**Movie 1.** Ciliary beating propels Trichoplax crawling on a glass coverslip.

**Movie 2.** Trichoplax move randomly on a clean glass substrate.

**Movie 3.** Trichoplax crawling movements are accompanied by body-shape changes.

**Movie 4.** Movie recording the movements of the animal analysed in Figure 2I-L, crawling on a glass coverslip on top of a plastic meshwork holder (background pattern).

**Movie 5.** Movie recording the movements of the animal analysed in Figure 2M-P, crawling on a glass coverslip on top of a plastic meshwork holder (background pattern).

**Movie 6.** Movie recording the movements of the animal analysed in Figure 3C,D, crawling on a glass coverslip on top of a plastic meshwork holder (background pattern). Poking with a round-tip scalpel modifies the direction of animal movement. 30sec after poking, the holder is transferred to cold acetone on dry ice.

**Movie 7.** Movie recording the movements of the animal analysed in Figure 3H-J, crawling on a glass coverslip on top of a plastic meshwork holder (background pattern). Bisecting the animal with a hand-held microscalpel results in the two animal halves moving away from each other. 30sec after cutting, the holder is transferred to cold acetone on dry ice.

**Movie 8.** Movie recording the movements of the animal analysed in Figure 3K-M, crawling on a glass coverslip on top of a plastic meshwork holder (background pattern). The animal is bisected with a hand-held microscalpel and 3-5sec after cutting the holder is transferred to cold acetone on dry ice.

**Movie 9.** Trichoplax incubated for 1hr in 50mM EGTA in FASW move slower than the controls (see Movie 2).

**Movie 10.** The speed of ciliary beating of Trichoplax incubated in 50mM EGTA is significantly decreased compared with controls (see Movie 1).

**Movie 11.** Movie recording the movements of the EGTA-treated animal analysed in Figure 4C-E, crawling on a glass coverslip on top of a plastic meshwork holder (background pattern).

**Movie 12.** Movie recording the movements of the EGTA-treated animal analysed in Figure 3I,J, crawling on a glass coverslip on top of a plastic meshwork holder (background pattern). Following bisection, the two animal halves fail to move away from each other. 30sec after cutting, the holder is transferred to cold acetone on dry ice.

**Movie 13.** Trichoplax incubated for 1hr in 75µM Verapamil in FASW move slower than the controls, but faster than the EGTA-treated animals (see Movies 2 and 9).

**Movie 14.** Movie recording the movements of the Verapamil-treated animal analysed in Figure 5C-E, crawling on a glass coverslip on top of a plastic meshwork holder (background pattern).

**Movie 15.** Movie recording the movements of the Verapamil-treated animal analysed in Figure 5I,J, crawling on a glass coverslip on top of a plastic meshwork holder (background pattern). Following bisection, the two animal halves fail to move away from each other. 30sec after cutting, the holder is transferred to cold acetone on dry ice.

## LEGENDS TO SUPPLEMENTARY FIGURES

**Figure S1.**
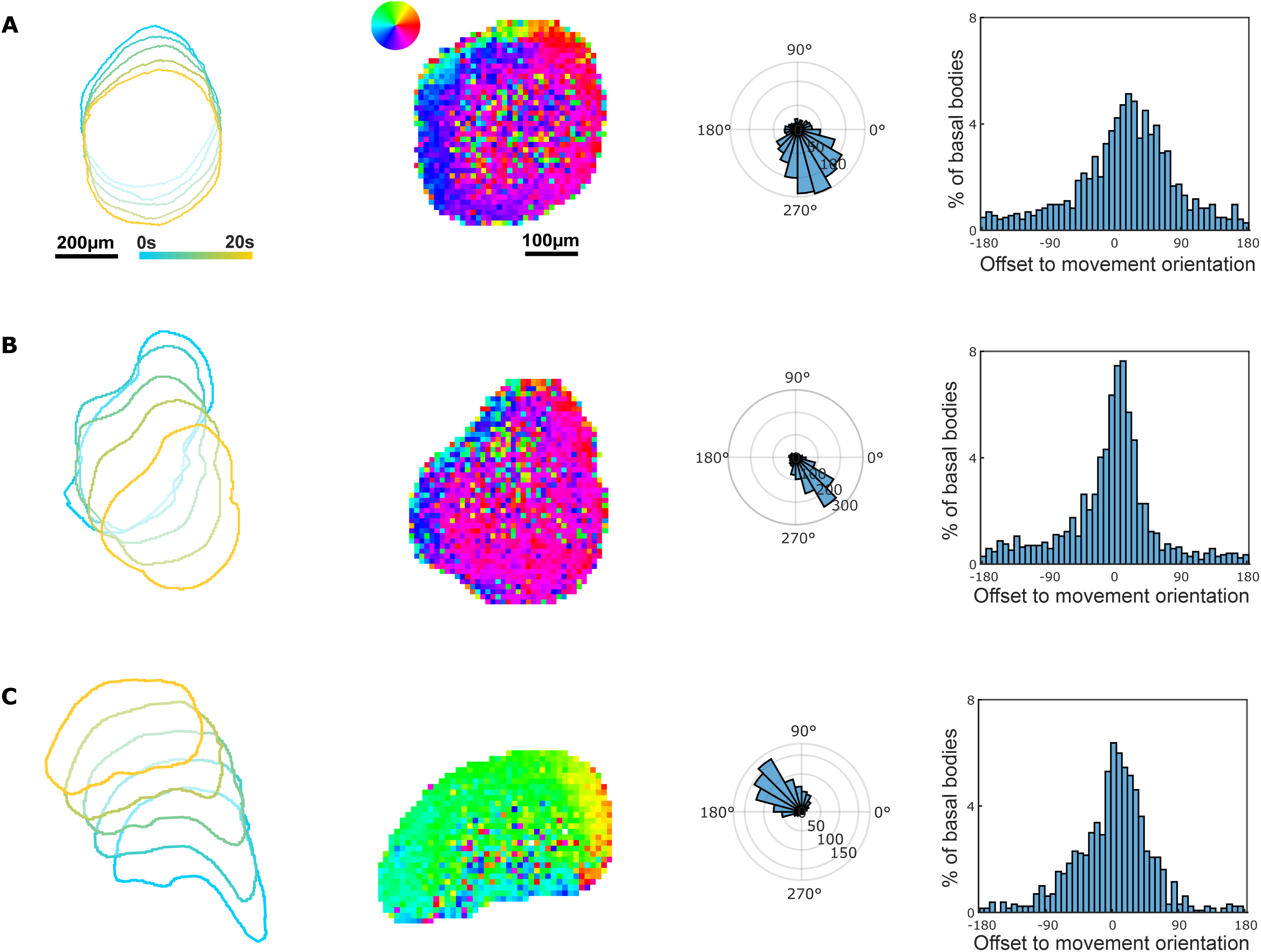
**A, B and C:** For three distinct Trichoplax, each row shows, from left to right: the moving body outlines, drawn at 5 second intervals; the colour-coded map of BB polarity; the BB polarity histogram; the offset of BB orientation respective to the direction of movement.

**Figure S2.**
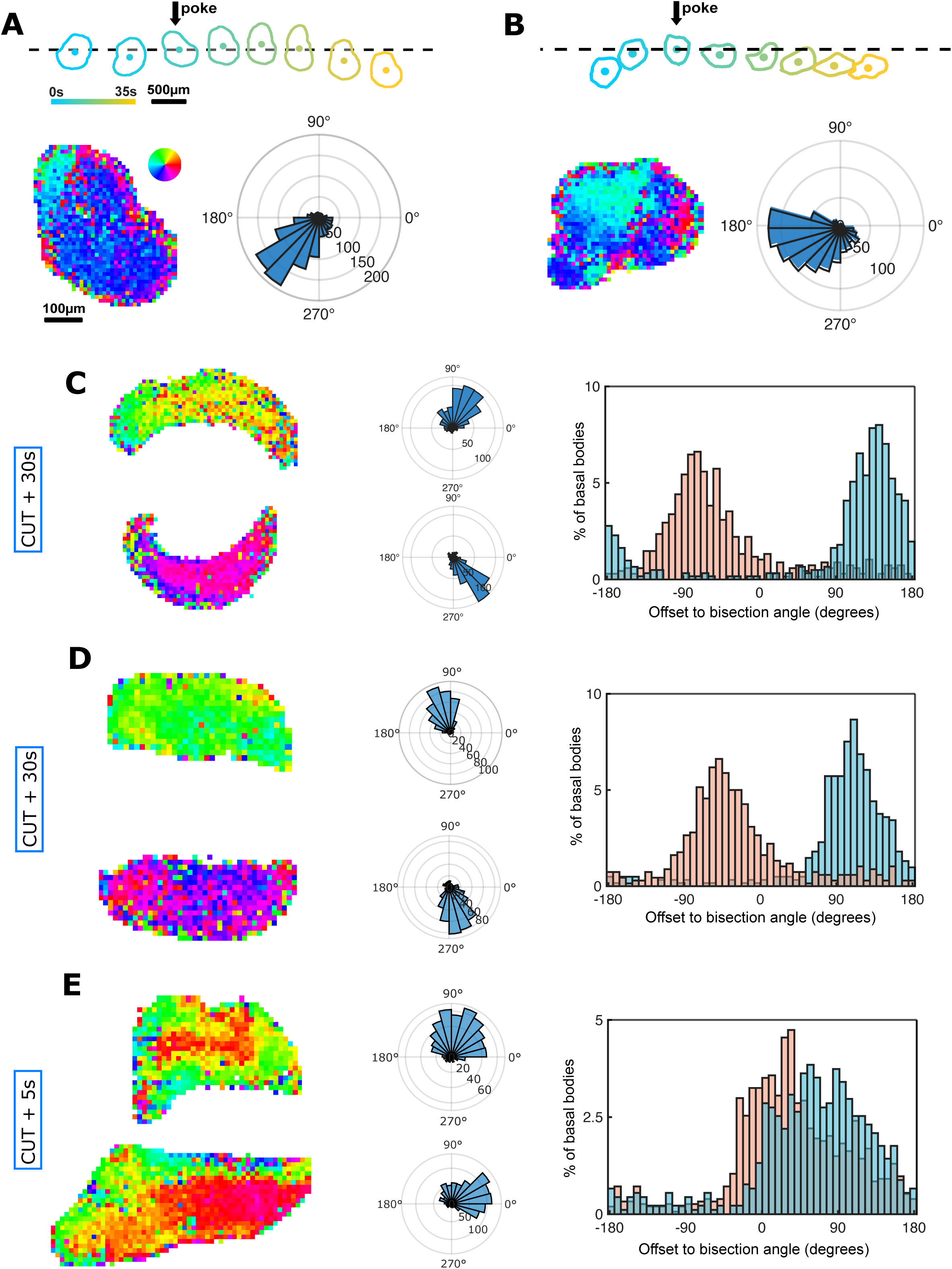
**A and B:** For two distinct Trichoplax, each panel shows: the moving body outlines, drawn at 5 second intervals, before and after poking (black arrow); the colour-coded map of BB polarity; the BB polarity histogram. **C and D:** For two distinct Trichoplax, fixed 30sec after bisection, each row shows, from left to right: the colour-coded map of BB polarity; the BB polarity histogram; the offset of BB orientation in the two halves respective to the cut. **E:** as in C and D, but for an animal fixed 3-5 seconds after bisection.

**Figure S3.**
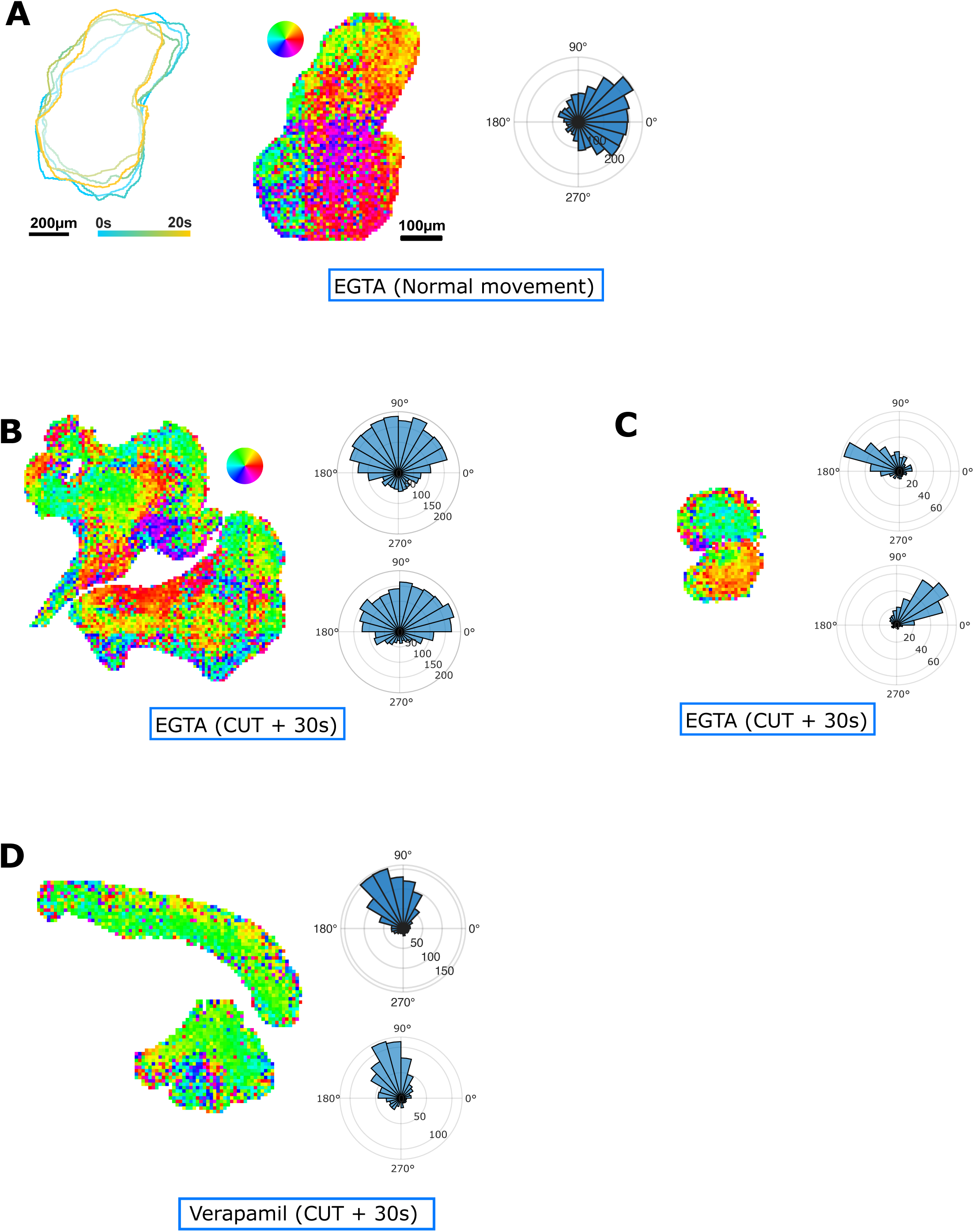
**A:** moving body outlines, drawn at 5 second intervals, colour-coded map of BB polarity and BB polarity histogram for an EGTA-treated Trichoplax. **B and C:** colour-coded map of BB polarity and BB polarity histogram for two EGTA-treated Trichoplax, fixed 30sec after bisection. **D:** as in B and C, but for a Verapamil-treated animal.

