## Supplementary figures and images for "Fast calcium-dependent reorientation of motile cilia basal bodies in the simple metazoan, Trichoplax"

### Supplementary Figures 1_3

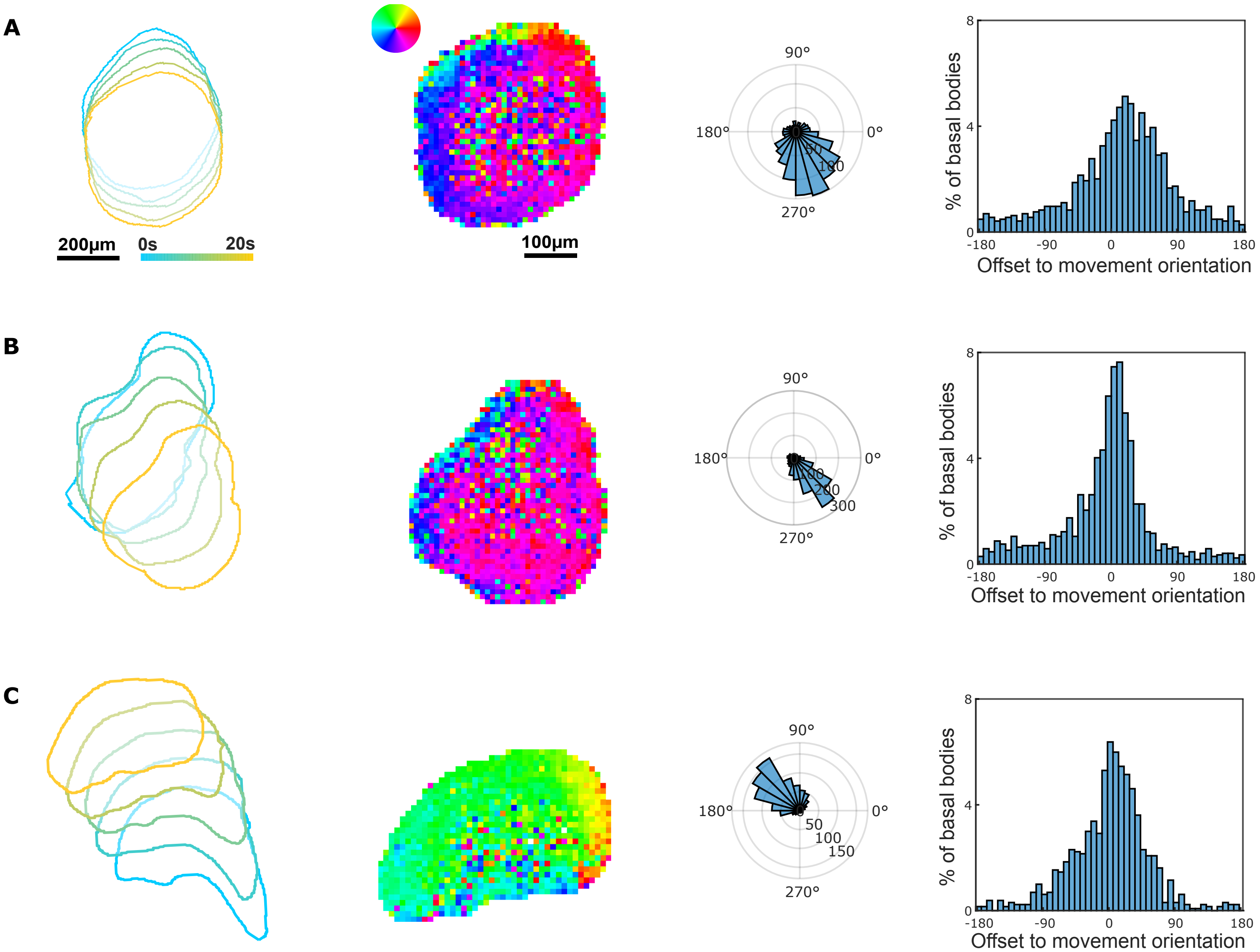


#### SUPPLEMENTARY FIGURE 1

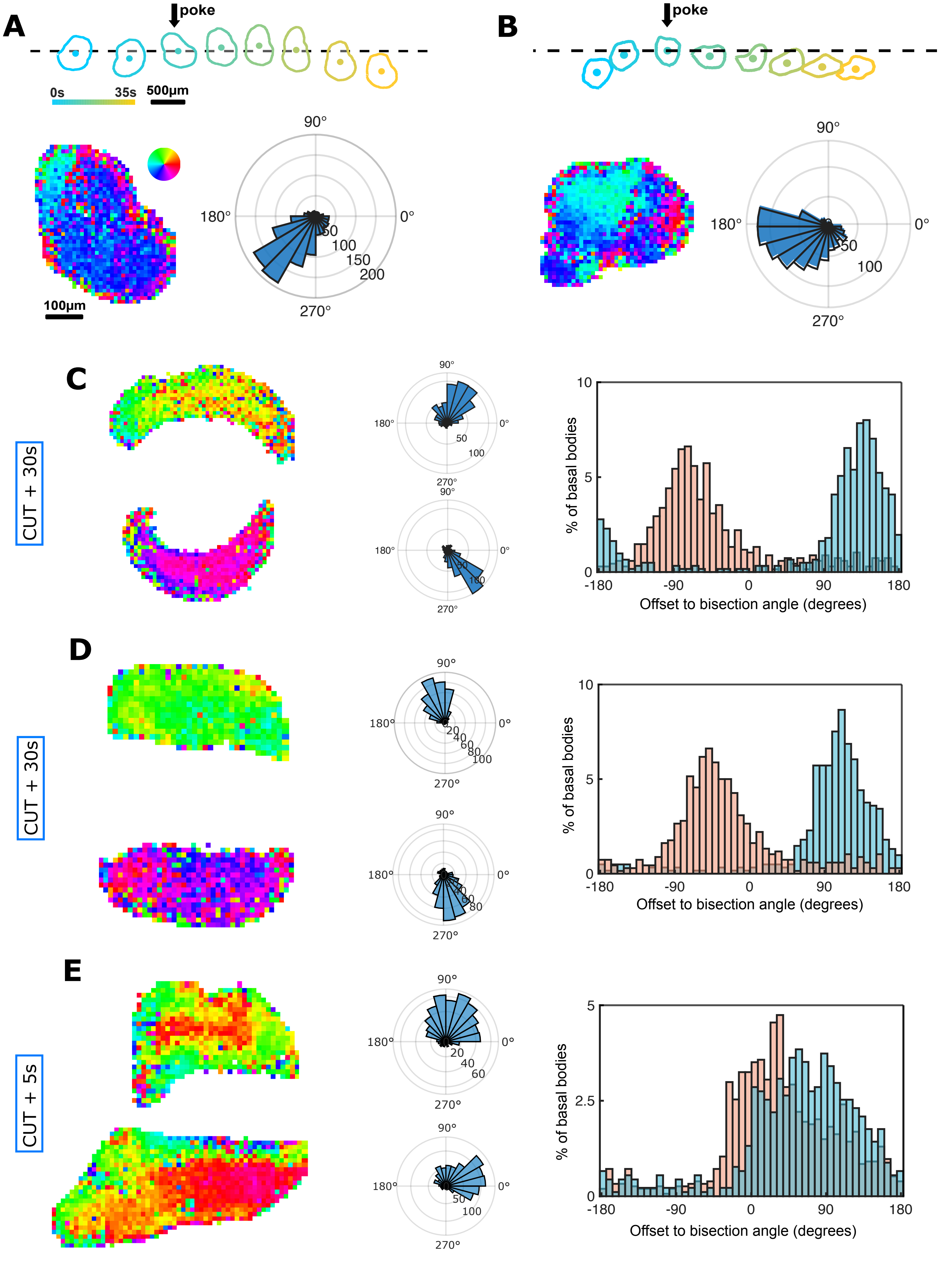


**SUPPLEMENTARY FIGURE 2**

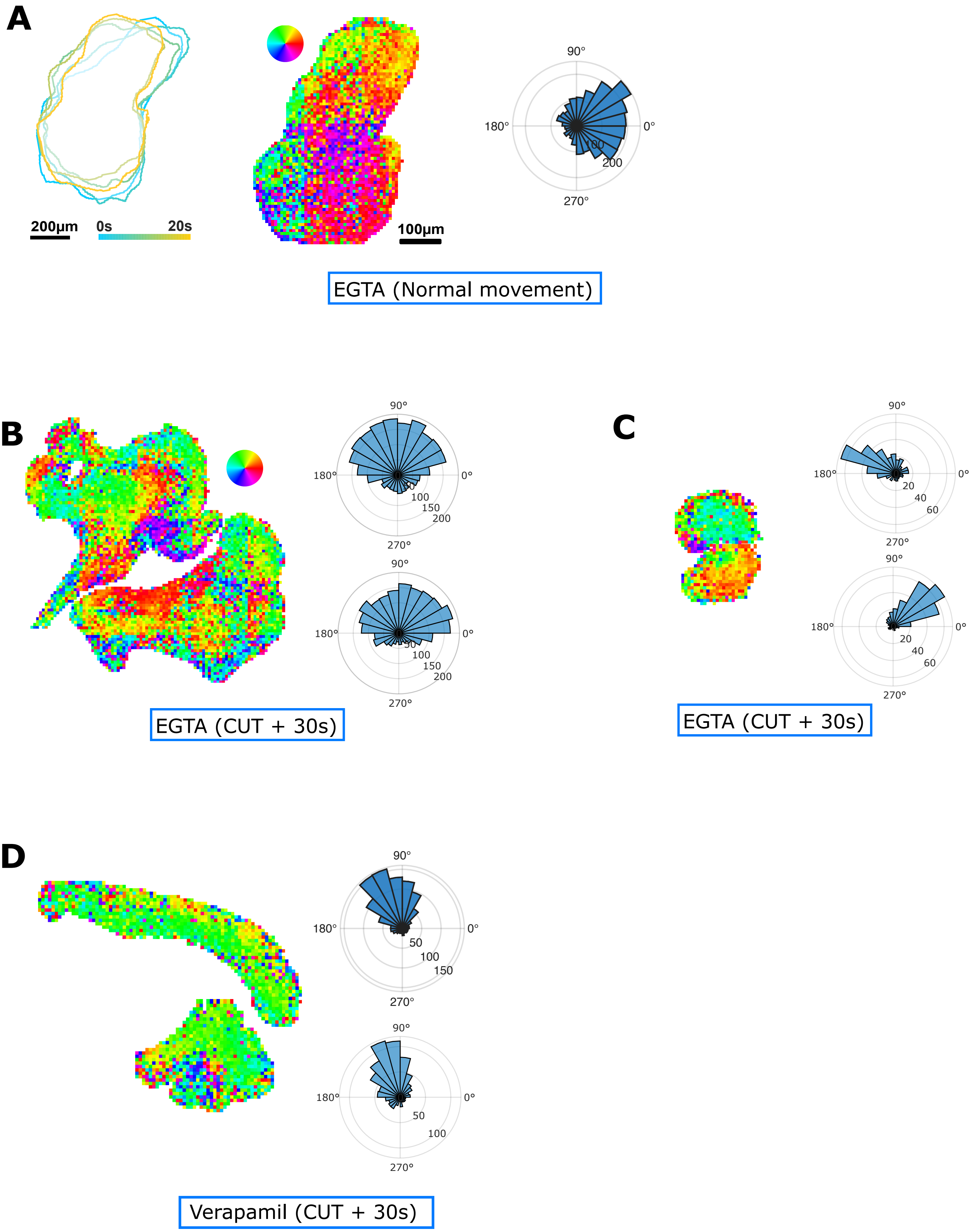


**SUPPLEMENTARY FIGURE 3**
